# Structure and function of BcpE2, the most promiscuous GH3-family beta-glucosidase for scavenging glucose from heterosides

**DOI:** 10.1101/2022.01.22.477351

**Authors:** Benoit Deflandre, Cédric Jadot, Sören Planckaert, Noémie Thiébaut, Nudzejma Stulanovic, Raphaël Herman, Bart Devreese, Frédéric Kerff, Sébastien Rigali

## Abstract

Cellulose being the most abundant polysaccharide on earth, beta-glucosidases hydrolyzing cello-oligosaccharides are key enzymes to fuel glycolysis in microorganisms developing on plant material. In *Streptomyces scabiei*, the causative agent of common scab in root and tuber crops, a genetic compensation phenomenon safeguards the loss of the gene encoding the cello-oligosaccharide hydrolase BglC by awakening the expression of alternative beta-glucosidases. Here we reveal that the BglC compensating enzyme BcpE2 is the GH3-family beta-glucosidase that displays the highest reported substrate promiscuity able to release the glucose moiety of all tested types of plant-derived heterosides (aryl β-glucosides, monolignol glucosides, cyanogenic glucosides, anthocyanosides, and coumarin heterosides). BcpE2 structure analysis highlighted a large cavity in the PA14 domain that covers the active site, and the high flexibility of this domain would allow proper adjustment of this cavity for disparate heterosides. The exceptional substrate promiscuity of BcpE2 provides microorganisms a versatile tool for scavenging glucose from plant-derived nutrients that widely vary in size and structure. Importantly, scopolin is the only substrate commonly hydrolyzed by both BglC and BcpE2 thereby generating the potent virulence inhibitor scopoletin. Next to fueling glycolysis, both enzymes thus also interfere with the plant defense mechanisms to fine-tune the strength of virulence.

## Introduction

The major source of soil organic carbon derives from plant senescence, with cellulose, xylan, starch, and lignin being the most abundant naturally-occurring carbon-containing polymers. Optimal colonization of carbon rich environments thus mainly relies on the ability of microorganisms to participate and feed on decomposing plant biomass. The same rationale applies to phytopathogenic bacteria for which the energy required for plant colonization must be supported by catabolic pathways biased for carbon sources utilization. Filamentous Gram-positive bacteria of the genus *Streptomyces* are well-known for their role in soil mineralization and their capability to consume diverse poly-, oligo-, and monosaccharides ^1–4^. Cellulose being the most abundant polysaccharide on earth, the acquisition of a complete cellulolytic system provides a competitive advantage in organic environments. However, as cellobiose is the main product resulting from cellulolysis ^5,6^, possessing the CebEFG-MsiK cello-oligosaccharide transporter ^7–9^ and the beta-glucosidase BglC ^10,11^ for their subsequent hydrolysis into glucose, would be a sufficient requirement for the survival of streptomycetes within a microbial community consuming plant material ^2^.

For *Streptomyces scabiei*, the bacterium responsible for common scab in root and tuber crops, cellobiose and cellotriose are of particular importance. Indeed, their import and subsequent catabolism do not only feed glycolysis with glucose, but they are also the environmental triggers of the onset of its pathogenic lifestyle ^7,12–17^. This tight link of cellulose byproduct utilization for both primary metabolism and the onset of virulence is perfectly highlighted by the phenotype of the null mutant of gene *scab57721* encoding the β-glucosidase BglC. Indeed, strain *S. scabiei* Δ*bglC* poorly grows when cultured with cellobiose or cellotriose provided as sole carbon sources, and also displays a hypervirulent phenotype due to the constitutive production of thaxtomin A, the main virulence determinant ^10^. Surprisingly, we recently showed that BglC is also able to remove the glucose moiety of the scopolin heteroside^18^ produced by plants under host colonization thereby generating scopoletin, a potent inhibitor of thaxtomin A production^19^. The hydrolysis of scopolin by BglC displays a substrate inhibition kinetic profile ^18^ that contrasts with the typical Michaelis–Menten saturation curve observed for the degradation of its natural substrates cellotriose and cellobiose^10^. At low concentration, the scopolin and cellobiose/cellotriose degradation would synergistically reduce thaxtomin A production and or feed glycolysis, while at higher concentrations the activities on both substrates would instead activate the biosynthesis of this main virulence factor. This enzyme has thus different kinetic properties on either the virulence elicitors emanating from the plant host or a molecule produced by the plant defense mechanisms, thereby occupying a key position to fine-tune the production of thaxtomin A^18^.

Surprisingly, we showed that the deletion of *bglC* in *S. scabiei* is not lethal when the mutant strain is inoculated on media with cellobiose supplied as unique carbon source^20^. The unexpected survival of the *bglC* null mutant in this culture condition is due to a genetic compensation phenomenon that awakens the expression of the gene *scab2391* encoding an alternative GH1-family β-glucosidase^20^. This sugar hydrolase was called BcpE1 (SCAB_2391), BcpE standing for <u>B</u>glC <u>c</u>om<u>p</u>ensating <u>e</u>nzyme, and is a paralogue of BglC, both enzymes having catalytic properties in the same order of magnitude towards cellobiose^20^. Interestingly, the genetic compensation phenomenon associated with the loss of *bglC* resulted in the overexpression of a second gene (*scab64101*) encoding a GH3-family β-glucosidase called BcpE2 ^20^. However, in contrast to BcpE1, BcpE2 cannot hydrolyze cellobiose^20^ and therefore the reason why the *bglC*-dependent mechanism of compensation selected the product of *scab64101* as alternative β-glucosidase remained unknown. If the role of BcpE2 is not to provide glucose from cellobiose or cellotriose, how could this enzyme compensate for the loss of function of BglC? In other words, how would *S. scabiei* and other streptomycetes benefit from the activation of BcpE2 in their environmental niche?

In this work we investigated through enzymatic, structural, and expression studies the function of BcpE2. Our results demonstrate that BcpE2 is able to release the glucose moiety of various types of plant-derived heterosides. Thanks to its exceptional substrate promiscuity, BcpE2 safeguards the feeding of glycolysis with glucose hydrolyzed from highly diverse carbon sources. Moreover, BcpE2 also degrades scopolin into scopoletin following a substrate inhibition profile, thereby also compensating for the loss of the function of BglC towards the host defense mechanism.

## Results

### Structure of BcpE2 of *Streptomyces scabiei*

We obtained the crystallographic structure of BcpE2 at 3.09 Å. The crystal belongs to the P3_1_21 space group with one molecule in the asymmetric unit. The BcpE2 structure is characterized by R_work_ and R_free_ values of 22.3% and 27% respectively (Table S1) and contains residues 10 to 822. Six regions could not be built because of a lack of electron density: the first nine amino acids (AA)-, the twelve C-terminal residues, which consist in a His_6_-Tag and a linker, and four loops (residues 70 to 73, 336 to 339, 436 to 443 and 531 to 536). BcpE2 is monomeric and is made of four domains (Figure 1a, Table S1): an N-terminal (β/α)_8_ TIM barrel domain (residues 10–306), an (α/β)_6_-sandwich domain (residues 316–398 and 551–650), a PA14 domain (residues 404–548), and a C-terminal fibronectin type III-like (fn3) domain (residues 711–821). In addition, a 60 AA linker is present between (α/β)_6_-sandwich and the fibronectin type III-like domains and mostly runs on the (β/α)_8_ TIM barrel domain. This architecture was identified in only 19 sequences among all 199 characterized prokaryotic and eukaryotic GH3s by mining the CAZy database. Only two structures, KmBglI from *Kluyveromyces marxianus* (PDB code 3AC0)^21^, and DesR from *Streptomyces venezuelae* (PDB code 4I3G) ^22^ were found with this architecture. While the (β/α)_8_ TIM barrel, the (α/β)_6_-sandwich, and the fn3 domains are very well superimposed, including the linker preceding the C-terminal domain (rmsd of 0.91 Å over 507 Cα for KmBglI and 0.90 Å over 504 Cα for DesR), the PA14 domain cannot be superimposed simultaneously with the three other domains (Figure 1bc). In BcpE2 and KmBglI, the orientation is roughly similar, and they are characterized by a rmsd of 2.6 Å over 81 Cα when superimposed independently. In DesR, the PA14 domain is 40 AA shorter than in BcpE2 and approximately perpendicular to the orientation in the latter. When superimposed independently, the rmsd between them is 4.7 Å over 88 Cα. The PA14 domain of BcpE2 is also characterized by B factor values significantly higher than the rest of the protein, particularly in the active site vicinity (Figure 1d), indicating a likely flexibility of this domain. This feature was also noted for KmBglI and potentially associated with substrate recognition ^21^.

**Figure 1.**
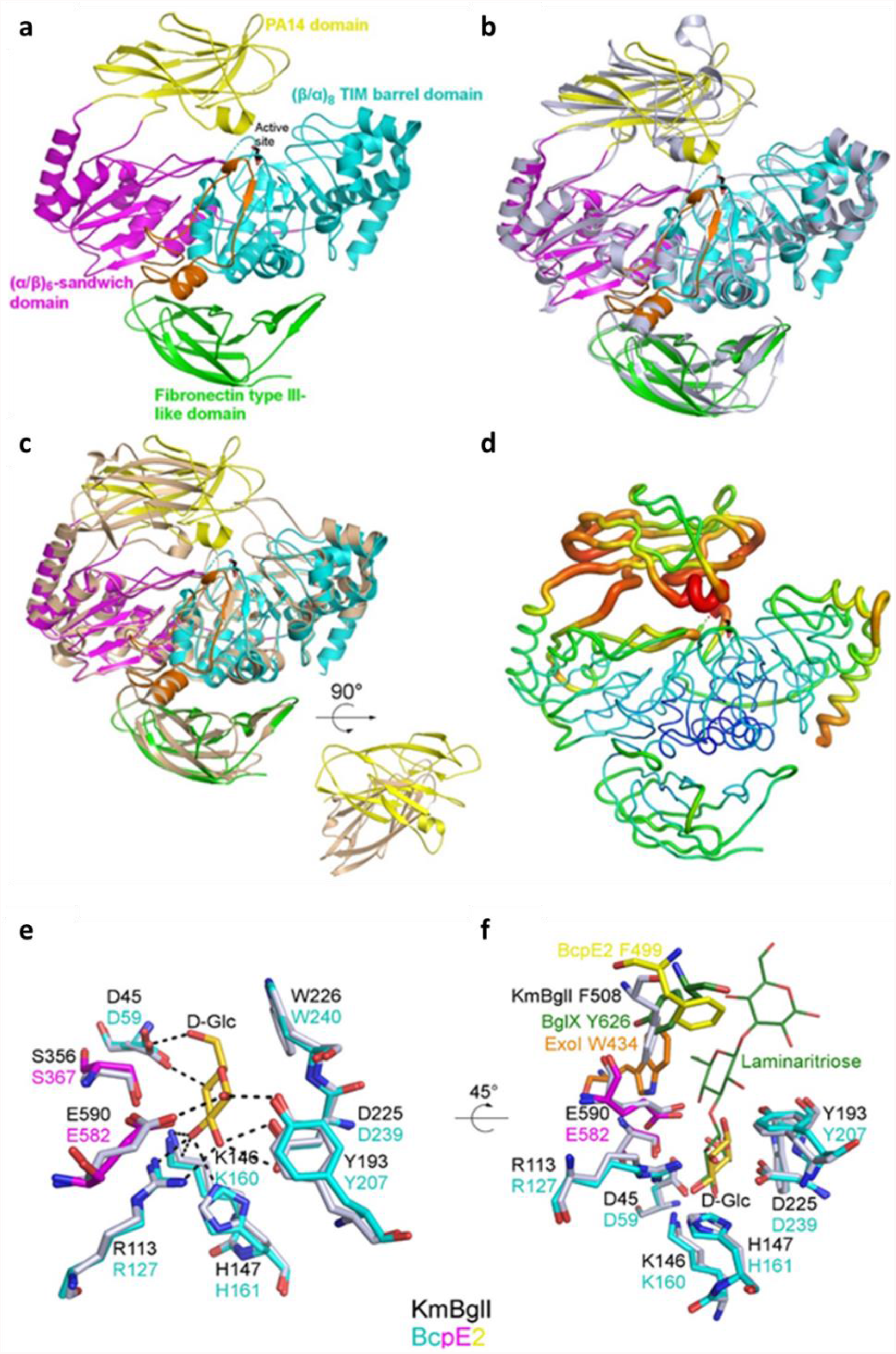
Overall fold and active site description of BcpE2 of *S. scabiei*. (**a**) Cartoon representation of the BcpE2 structure. The glycerol molecule in the active site (black sticks) results from the cryo-protectant solution used for freezing the crystal. (**b**) Superimposition of BcpE2 and KmBglI (light blue). (**c**) Superimposition of BcpE2 and DesR (light orange). Their PA14 are also shown with a 90° rotation to highlight the different orientations. (**d**) Ribbon representation of BcpE2. The ribbon radius is proportional to the mean B-factor value of the residues and a rainbow coloring scheme (blue low B factor value to red high B-factor value). (**e**) Superimposition of the catalytic site of BcpE2 (residues are colored by domain with the same coloring scheme as in (b) with KmBglI (grey) in complex with D-Glucose (D-Glc in gold) in subsite -1. Hydrogen bonds with D-Glc are shown as black dashed lines. (**f**) Same as (e) with a 45° rotation and the addition of ExoI (orange) and the BglX:Laminaritriose complex (green) superimposed. The aromatic residues of subsite +1 of KmBglI and BglX are also shown as sticks, as well as their likely equivalent in ExoI and BcpE2. The position of F499 in BcpE2 (yellow sticks) comes from the PA14 domain superimposed independently on the PA14 domain of KmBglI.

### The catalytic site of BcpE2

Both KmBglI and DesR have been crystallized with a β-D-glucose molecule present in their active site (−1 subsite) ^21–23^. This molecule corresponds to the buried end of the substrate, and, in KmBglI, it is found in a pocket made of 9 residues (7 of them being involved in hydrogen bonds) belonging to the (α/β)_8_ TIM barrel and the (α/β)_6_-sandwich domains. These residues, which include the catalytic glutamate and aspartate responsible for the two-step double displacement mechanism ^24^, are strictly conserved with a very similar orientation in DesR and BcpE2 (rmsd calculated for all non-hydrogen atoms of 0.48 Å and 0.60 Å respectively; Figure 1e). Figure S1 shows the structure-based sequence alignment of BcpE2 with homologous and biochemically characterized proteins (Table S2) that either display the highest structural or primary sequence similarity. The two catalytic residues – Asp239 and Glu582 – are strictly conserved in all homologous GH3s (Figure S1). These residues provide an ideal geometry to stabilize the five hydroxyl groups of glucose with at least one hydrogen bond and facilitate an efficient hydrolysis. The active site of BcpE2 also includes Asp59, Arg127, and three additional aromatic AAs, namely Tyr207, Trp240, and Phe499 which are generally conserved in the closest GH3-family enzymes (Figure S1).

In KmBglI, Phe508, which belongs to the PA14 domain, has been identified by site directed mutagenesis as important for substrate hydrolysis and is part of the subsite (+1)^25^. In BcpE2, Phe499 is equivalent when the two PA14 domains are superimposed independently (Cα 2 Å apart) but is shifted toward the position that the substrate would likely occupy when the entire structures are superimposed (Cα 5 Å apart). This positioning could be the result of the crystal packing and/or due to the flexibility observed. An aromatic residue at this position seems to be a common feature in GH3 enzymes but can come from different structural elements (Figure S1). For example, in the ExoI enzyme form barley^26^, it is located on a loop of the (α/β)_6_-sandwich domain, and in the dimer forming BglX from *Pseudomonas aeruginosa* ^24^, it comes from an extended loop of the second molecule of the dimer (Figure 1f). In BcpE2, the high B factor values observed in this region and the likely related flexibility make it difficult to extract additional information about substrate specificity in the (+1) subsite at this stage.

### Seeking for the natural substrate(s) of BcpE2

Earlier work showed that BcpE2 displayed a strong hydrolytic activity on the synthetic chromogenic substrate pNPβG^20^, suggesting that the enzyme should target carbohydrates with a terminal glucose attached by a β-1,4 linkage. According to KEGG pathway, BcpE2 of *S. scabiei* 87-22 is suggested as candidate beta-glucosidase possibly involved in cyanoamino acid metabolism (https://www.genome.jp/kegg-bin/show_pathway?scb00460+SCAB_64101). Two cyanogenic glucosides were thus selected as possible targets of BcpE2, *i.e*., amygdalin and linamarin (Table 1). To help identifying other putative substrate(s) that could be hydrolyzed by BcpE2, we generated a phylogenetic tree with BcpE2 of *S. scabiei* and the full-length sequences of the fourteen closest characterized bacterial GH3-family β-glucosidases, and with five other characterized bacterial GH3s with lower overall identity but with high query coverage and containing the PA14 domain (Figure S2).

**Table 1.** Overview of the activity of BcpE2 and BglC on selected disaccharides and heterosides.

| Substrate | Category | TLC assays |  | Kinetic parameters (BcpE2) |  |  |  |
| --- | --- | --- | --- | --- | --- | --- | --- |
|  |  | BcpE2 | BglC | K<sub>m</sub> (mM) | k<sub>cat</sub> (s<sup>-1</sup>) | K<sub>m</sub>/k<sub>cat</sub> (mM<sup>-1</sup>.s<sup>-1</sup>) | K<sub>i</sub> (mM) |
| **Cellobiose** | Disaccharides (β-1,4 glucose) | - | + | /<sup>20</sup> | /<sup>20</sup> | /<sup>20</sup> | /<sup>20</sup> |
| **Xylobiose** | Disaccharides (β-1,4 xylose) | - | - | NT | NT | NT | NT |
| **Laminaribiose** | Disaccharides (β-1,3 glucose) | ± | + | 6.557<br>± 1.410 | 8.74<br>± 1.02 | **1.33** | NA |
| **Gentiobiose** | Disaccharides (β-1,6 glucose) | ± / - | ± / - | NT | NT | NT | NT |
| **Salicin** | Aryl-β-glucosides | + | ± | 0.142<br>± 0.030 | 53.20<br>± 2.37 | **375.71** | NA |
| **Arbutin** | Aryl-β-glucosides | + | ± / - | 0.367<br>± 0.080 | 43.17<br>± 2.50 | **117.49** | NA |
| ***p*-coumaryl alcohol**<br>**4-O-glucoside** | Monolignol<br>glucosides | + | ± | 0.236 | 77.15 | **326.49** | 0.593 |
| **Syringin** | Monolignol<br>glucosides | + | - | 0.608<br>± 0.166 | 26.43<br>± 3.07 | **43.46** | NA |
| **Coniferin** | Monolignol<br>glucosides | + | ± | 0.151<br>± 0.043 | 83.30<br>± 5.90 | **551.29** | NA |
| **Cyanin (Cyanidin-**<br>**3,5-di-O-glucoside)** | Antho-<br>cyanosides | + | - | NT | NT | NT | NT |
| **Esculin** | Coumarin<br>heterosides | + | ± | NT | NT | NT | NT |
| **Scopolin** | Coumarin<br>heterosides | + | + | 0.356 | 191.90 | **539.65** | 0.152 |
| **Linamarin** | Cyanogenic<br>glycosides | + | - | NT | NT | NT | NT |
| **Amygdalin** | Cyanogenic<br>glycosides | ± | - | NT | NT | NT | NT |
| **4-MUG** | Aryl-β-<br>glucosides | + | + | NT | NT | NT | NT |

According to these *in silico* analyses combined to a literature survey, 14 candidate natural substrates were selected for BcpE2 (Table 1). Most of them are β-1,4 linked heterosides commonly found in plants, *i*.*e*., compounds with glucose (or another carbohydrate moiety) linked by a glycosidic bond to an aglycone. They belong to different types of plant heterosides with extremely variable aglycone moieties, i.e., i) aryl-β-glucosides (arbutin, salicin), ii) monolignol glucosides (p-coumaryl alcohol 4-O-glucoside, syringin, coniferin), iii) anthocyanosides (cyanin), iv) coumarin heterosides (esculin, scopolin), and cyanogenic glycosides (linamarin, amygdalin).

The candidate substrates listed in Table 1 were tested to determine the ability of BcpE2 to release their glycosidic moiety (Figure 2) β-glucosidase BglC was also included in our enzymatic assays to compare the respective substrate specificities of each enzyme. Both pure six histidine-tagged enzymes were incubated with the candidate substrates and reactions were conducted at their optimal pH (7.5) and temperature (40°C) (optima deduced from results presented in Figure S3 for BcpE2 and described in ^10^ for BglC). Reaction samples were spotted on thin layer chromatography (TLC) plates and migrated in an elution chamber to separate glucose (or other saccharides) from the remainder moieties of the substrate (Figure 2a).

**Figure 2.**
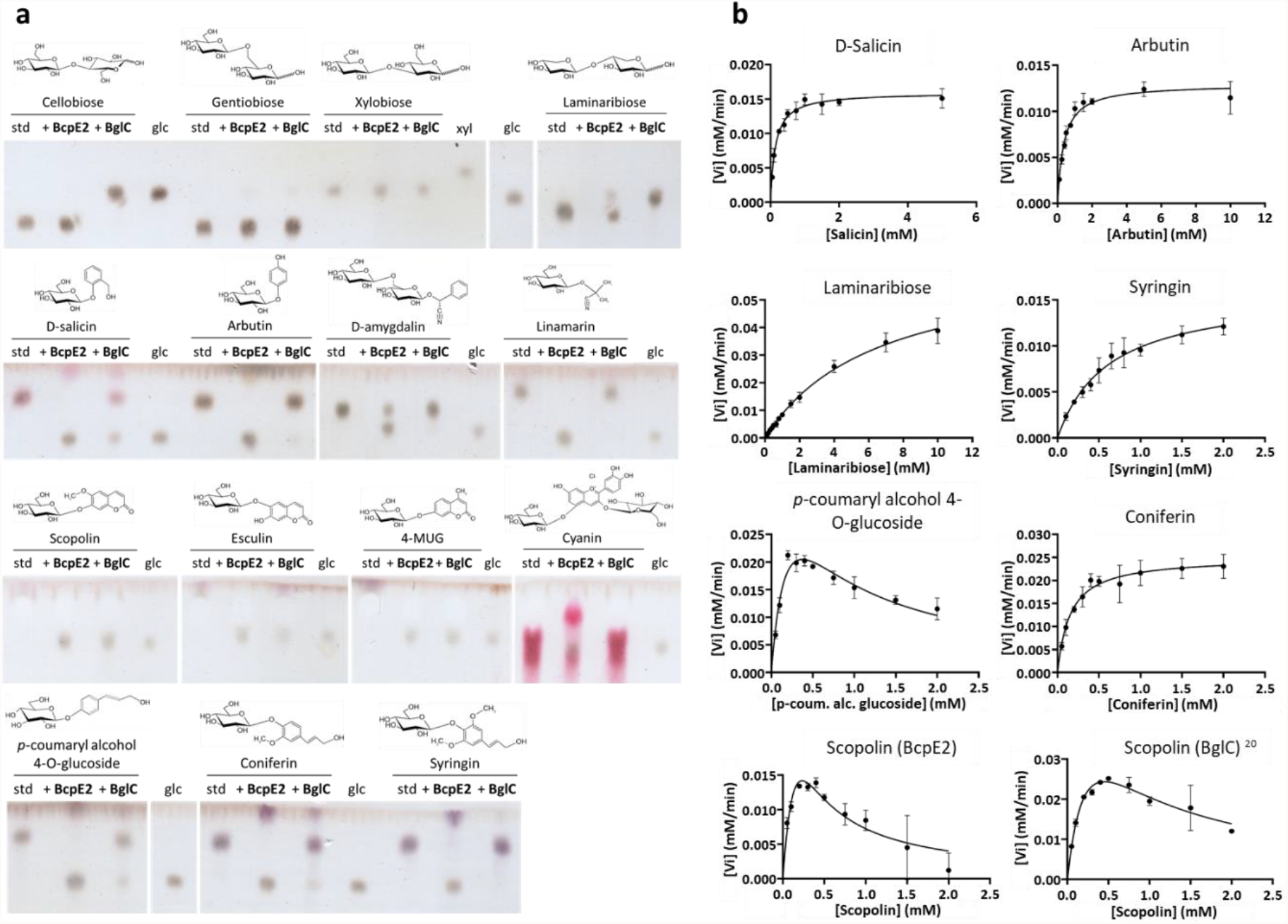
Substrate specificities of BcpE2 and BglC. (**a**) TLC plates revealing the release of glucose (glc) after incubation of a variety of substrates (5 mM) with BcpE2 or BglC (1 µM) compared to the intact substrate (standard (std)). The chemical structure is displayed above each substrate. (**b**) Non-linear regressions of the kinetic analyses of BcpE2 towards seven substrates and one for BglC. Plots of the initial velocity (Vi, mM/min) estimated by the rate of glucose released by the enzyme as a function of substrate concentrations (in mM). Individual values were entered into the GraphPad Prism software (9.2.0) which fitted the data to the Henri-Michaelis-Menten model by a non-linear regression. In the case of a decrease of the Vi at high substrate concentrations, the data were fitted to the Substrate Inhibition model. Error bars display the standard deviation values determined for the Vi by three replicates at each substrate concentration.

Cellobiose was first tested as positive and negative control substrate for BglC and BcpE2, respectively. Indeed, as shown in Figure 2a, glucose is only released from cellobiose when this substrate is incubated with BglC. Other D-glucose disaccharides were tested, namely gentiobiose (D-glucose linked in β(1→6)), and laminaribiose (D-glucose linked in β(1→3)). Surprisingly, BglC could efficiently degrade laminaribiose whereas BcpE2 could only partially degrade this substrate. Both enzymes were equally inefficient on gentiobiose where barely perceptible amounts of glucose were released (Figure 2a). Neither BglC nor BcpE2 was active on xylobiose suggesting that these enzymes cannot properly target D-xylose saccharides.

Strikingly, BcpE2 revealed to be active on all tested heterosides (Figure 2a). Complete hydrolysis was observed for i) the two aryl-β-glucosides salicin and arbutin, ii) the cyanogenic glucoside linamarin, iii) the pink/purple anthocyanoside Cyanidin-3,5-di-O-glucoside chloride (Cyanin), iv) all three monolignol glucosides syringin, coniferin and *p*-coumaryl alcohol 4-O-glucoside, v) the coumarin heteroside esculin, and vi) the synthetic substrate 4-MUG (Figure 2a). Significant yet incomplete hydrolysis by BcpE2 was also observed for the cyanogenic glucoside amygdalin. In contrast, BglC was inactive on most tested heterosides except the synthetic substrate 4-MUG, and scopolin as previously described^18^. Only partial substrate hydrolysis by BglC could be observed for esculin, coniferin, *p*-coumaryl alcohol 4-O-glucoside, and salicin (Figure 2a). Overall, we observed that the substrate specificity of BcpE2 is broad and often complementary to that of BglC (Table 1). Surprisingly and despite an extensive variability in the aglycone parts of the tested compounds, BcpE2 managed to generate glucose from all the heteroside substrates considered in this study.

After determining the best candidate substrates of BcpE2 by preliminary enzymatic assays on TLC, a subset of them was used to evaluate the kinetic parameters of BcpE2. The values of the Michaelis constant (K_m_), catalytic rate constant (or turnover; k_cat_) and catalytic efficiency (k_cat_/K_m_) were determined for BcpE2 towards the seven following substrates: salicin, arbutin, laminaribiose, syringin, *p*-coumaryl alcohol glucoside, coniferin, and scopolin (Table 1) based on the non-linear regressions displayed in Figure 2b.

The best affinity of BcpE2 was – as indicated by the lowest K_m_ values – observed towards salicin and coniferin with a K_m_ of about 0.15 mM. The other substrates displayed values on the same order of magnitude, except laminaribiose for which the 6.557 mM estimated K_m_ value indicates a low affinity of the β-glucosidase for this substrate (as already anticipated from assays on TLC plates Figure 2a). Regarding the turnover parameter, the monolignol glucoside coniferin was the most efficiently hydrolyzed substrate with a k_cat_ value of 83.3 s^-1^. This heteroside thus had the highest catalytic efficiency at about 550 mM^-1^.s^-1^, surpassing salicin and *p*-coumaryl alcohol glucoside by some margin, and is thereby the best reported substrate for BcpE2.

While most of the tested substrates exhibited conventional Henri-Michaelis-Menten behavior upon hydrolysis by BcpE2, two heterosides revealed substrate inhibition, namely the *p*-coumaryl alcohol glucoside and scopolin (Figure 2b). Indeed, the monolignol glucoside *p*-coumaryl alcohol glucoside showed a decrease in the initial velocity at high substrate concentration. The calculated inhibition constant (K_i_) value for *p*-coumaryl alcohol glucoside was estimated at about 0.6 mM and the theoretical K_m_ and k_cat_ values without the inhibition phenomenon would be of 0.236 mM and 77.15 s^-1^, respectively. This suggests that this monolignol glucoside is also a good substrate for BcpE2, yet at relatively low concentrations. Interestingly, scopolin is the only natural substrate to be hydrolyzed by both BcpE2 and BglC in the TLC experiment (Figure 2a). We therefore decided to evaluate their respective kinetic parameters to evaluate how efficiently they degrade this substrate. In order to obtain similar initial velocity values at low substrate concentrations, a concentration 20-times higher of BglC compared to BcpE2 was required, suggesting a much better k_cat_ for the latter. Also, as indicated by the aspect of the non-linear regressions in Figure 2b, both enzymes are subjected to substrate inhibition. However, the K_i_ value for BcpE2 appears to be lower thus indicating a stronger inhibition compared to BglC (Table 1).

### Production of BcpE2 is induced by the aryl-β-glucoside salicin

To validate the role of BcpE2 in heteroside degradation *in vivo*, it is mandatory to show that BcpE2 is produced when *S. scabiei* encounters these types of molecules in its environment. From all tested substrates that are best hydrolyzed by BcpE2, we chose salicin as a putative natural elicitor of BcpE2 production (also due to its availability in terms of cost and quantity) for our *in vivo* production assays. *S. scabiei* was cultivated under conditions that allow BglC production (minimal medium containing cellobiose) and/or with salicin as putative trigger for BcpE2 production. The different intracellular crude extracts were separated by anion exchange chromatography and the fractions obtained were first tested against pNPβG as substrate to detect those containing β-glucosidases, and then subjected to targeted proteomics for the identification and quantification of BglC and BcpE2. The semi-quantitative abundances of BglC and BcpE2 under the three tested culture conditions are presented in Figure 3.

**Figure 3.**
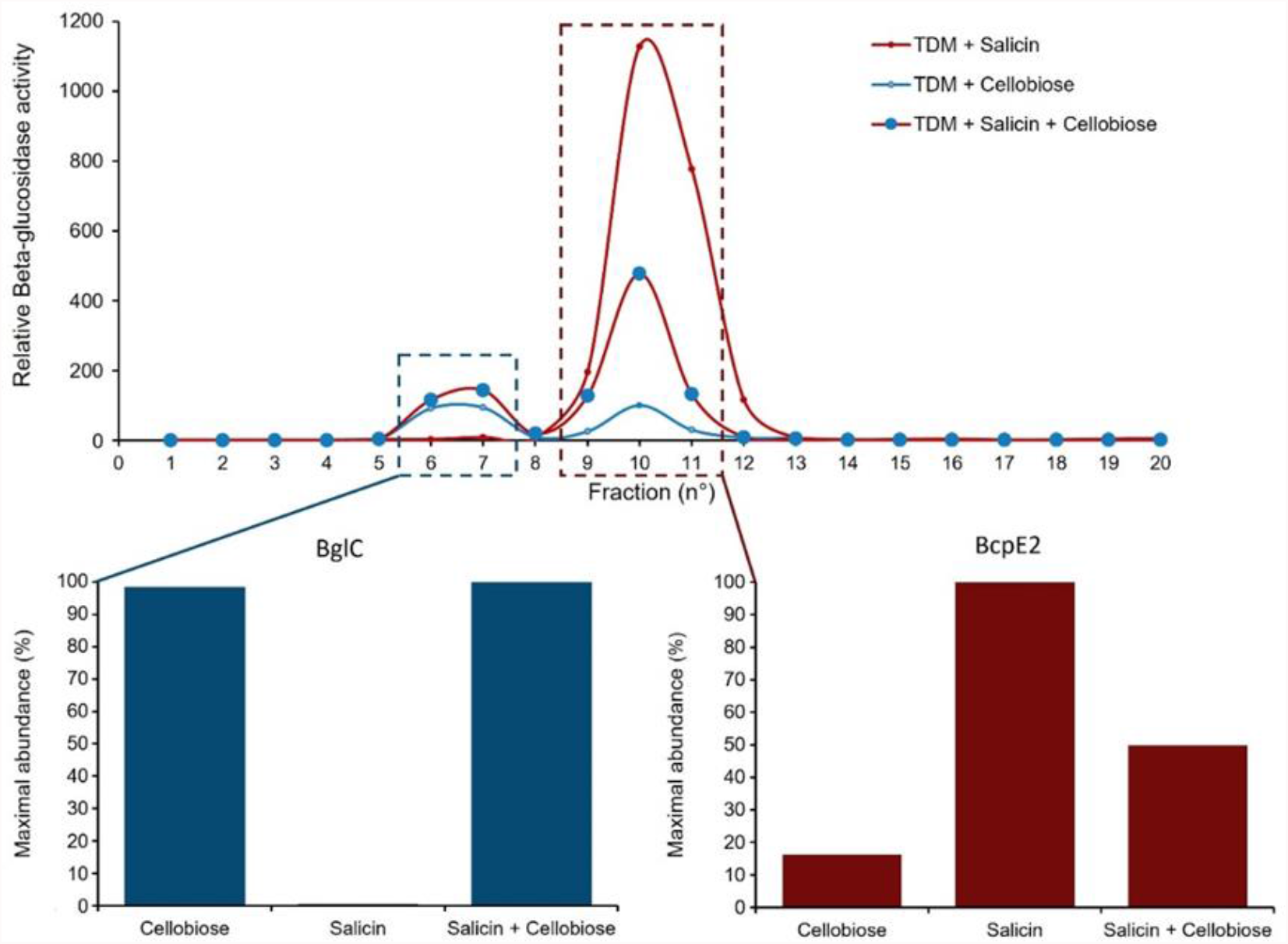
Induction of the respective production of BglC and BcpE2 by cellobiose and salicin. (Top pannel) Relative β-glucosidase activity in anion exchange chromatography fractions obtained from the full protein extracts of *S. scabiei* cultured in TDM medium supplemented with salicin (red trait), cellobiose (blue trait), or both substrates (red trait with blue circles). (Bottom pannel) Relative abundance of BglC in the first active peak (fractions 6-7) and of BcpE2 in the second active peak (fractions 9-11) determined by targeted proteomics (LC-MRM (Liquid chromatography multiple reaction monitoring) after tryptic digestion of the protein fractions). In each culture condition, the relative protein abundance was reported to the maximal abundance measured for the given protein.

As previously reported, the production of BglC is triggered by the presence of cellobiose. Salicin was neither able to induce (when provided as unique carbon source) nor repress (when in combination with cellobiose) the production of BglC suggesting that the expression of *bglC* is not under the control of this aryl β-glucoside. By contrast, BcpE2 was instead maximally produced when salicin was supplied as unique carbon source and the supply of cellobiose in addition to salicin reduced BcpE2 production to half of this level. When salicin was not supplied in the culture media, the production levels of BcpE2 dropped to about 16% of its maximal production level. Our results show that the production of BcpE2 is indeed triggered upon sensing the presence of salicin, one of the substrates for which the enzyme displayed the most efficient catalytic properties.

## Discussion

In this work, we report the structural and biochemical characterization of BcpE2, a GH3-family β-glucosidase of the common scab phytopathogen *S. scabiei*. This protein displays low similarity compared to biochemically characterized enzymes of this family, indicating that BcpE2 could have novel functional specificities. The crystal structure of BcpE2 revealed the presence of four domains – including a rather uncommon PA14 domain predicted to be involved in substrate specificity – organized around a catalytic pocket, which can accommodate D-Glucose as buried residue. BcpE2 was highly active against a wide variety of plant heterosides mostly containing glucose as carbohydrate residue, with salicin and coniferin as the most efficiently hydrolyzed substrates.

The BcpE2 structure reveals two features likely contributing to its broad substrate specificity, *e*., i) a wide cavity in the PA14 domain connected to the D-glucose specific subsite (−1), and the high flexibility of this domain, especially the structure elements defining this cavity (Figure 4). Three of the four loops not fully defined in the electron density are indeed adjacent to the cavity, which is also surrounded by the residues with the highest B factors. This will therefore provide sufficient plasticity to accommodate the various substrates and an easy access to efficiently load the substrates and expel the products of the hydrolysis. Except for the aromatic feature of Phe499, which seems to be a characteristic feature of GH3 enzymes, the residues of PA14 defining the cavity are not conserved even among closely related proteins (Figure S1). This could indicate that the main purpose of the cavity would be to hold the substrate shielded in the active site just long time enough for hydrolysis to take place, without contributing to the specificity except for setting a size limit.

**Figure 4.**
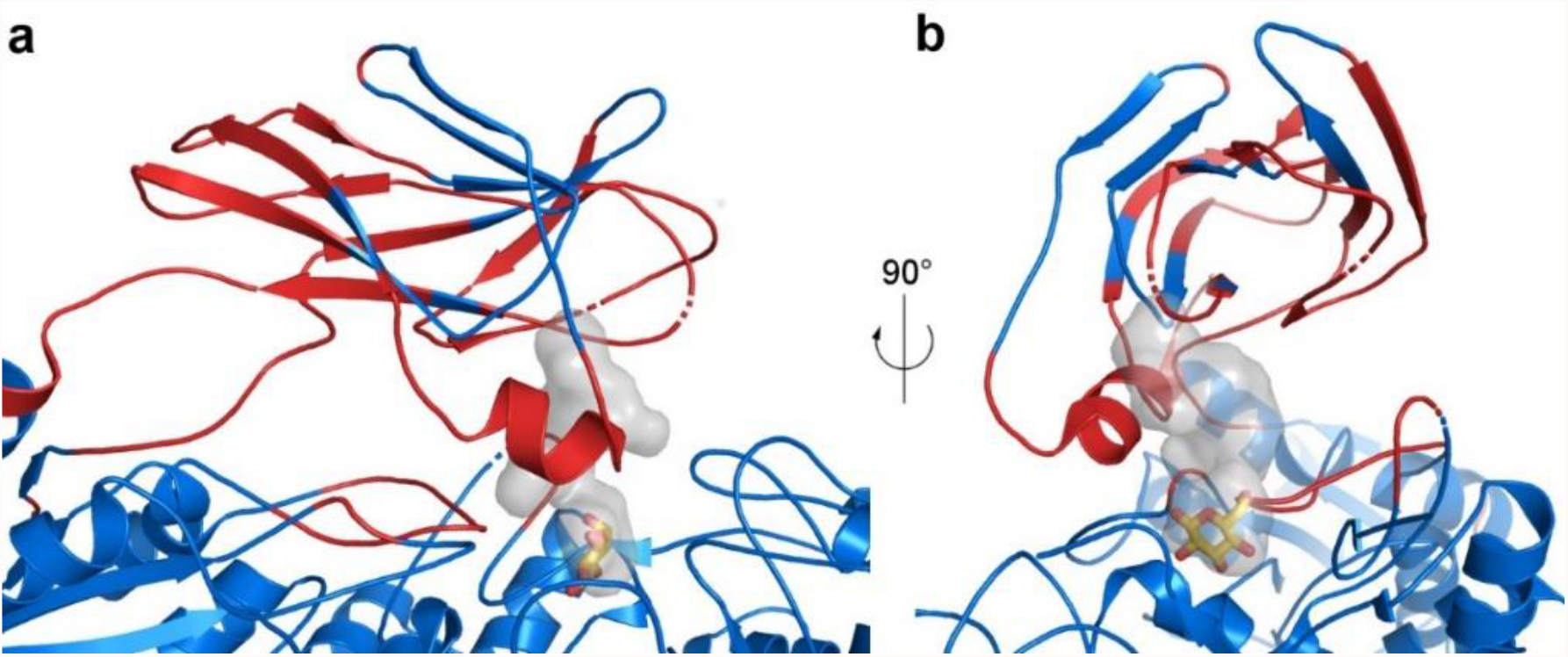
Active site flexibility of BcpE2. (**a**) Cartoon representation of the BcpE2 active site and the PA14 domain. Loops with missing amino acids are shown as dashed lines. Residues with a B factor of the Cα above 100 Å^2^ are in red and the others in blue. The active site pocket is represented with a transparent surface with the D-glucose molecule in the (+1) subsite from the superimposed KmBglII structure as yellow sticks. (**b**) Same as in (a) with a 90° rotation.

At this stage, the perhaps most difficult question to answer is “which heteroside could escape hydrolysis by BcpE2”, in other words, *to what extent can BcpE2 tolerate substrate promiscuity?* Glycosylated phytochemicals include phenylpropanoids, cyanogenic glucosides, coumarin heterosides, quinones, mono-or triterpenes, polyphenols, flavonoids (anthocyanosides, flavanols, isoflavonoids, flavonols and flavones), monolignols, amongst many others. Assessing the efficiency of BcpE2 to hydrolyze other substrates with an even wider spectrum of aglycone moieties will likely reveal the extent of its catalytic potential. A single heteroside, amygdalin, was not thoroughly hydrolyzed by BcpE2 (see TLC assay, Figure 2a), but it is also the only tested compound bearing gentiobiose instead of a glucose molecule as glycone. The gentiobiose disaccharide was poorly degraded by BcpE2 and it was thus not surprising that amygdalin was not thoroughly hydrolyzed. Despite this, the presence of the aromatic residue in the structure of this cyanogenic glucoside appears to enhance the activity of BcpE2 when compared to the hydrolysis of gentiobiose alone. The presence of at least one aromatic cycle in the chemical structure of the heterosides degraded by BcpE2 appears to be a common feature except for linamarin (Figure 2a). The fact that this cyanogenic glucoside is also efficiently hydrolyzed by BcpE2 suggests that the presence of an aromatic residue in the aglycone is not a mandatory feature to be accommodated as a substrate.

### Why is BcpE2 selected for compensating the loss of BglC?

As cello-oligosaccharides are not natural substrates of BcpE2, the reason why the *bglC*-dependent mechanism of genetic compensation selected the product of *scab64101* as alternative β-glucosidase was a complete mystery^20^. In light of the discovery of the substrate and enzymatic specificities of BcpE2, it is now easier to understand why the product of this gene was selected to compensate for the loss of *bglC*/BglC. Indeed, we now know that BcpE2 can compensate the impaired cello-oligosaccharide consumption by feeding glycolysis with glucose hydrolyzed from a multitude of plant-derived heterosides including monolignol glycosides which are some of the most ubiquitous molecules (Figure 5). BcpE2 would therefore act as a glucose scavenging enzyme, able to provide the most readily metabolized carbon source from a plethora of compounds that *S. scabiei* would encounter during host colonization. For soil-dwelling saprophytic streptomycetes, BcpE2 would be equally important as most of the organic carbon are molecules generated from plant senescence. Indeed, genome mining of streptomycetes revealed that most of them possess an orthologue of BcpE2. It is also striking that BglC and BcpE2 possess complementary substrate ranges on the tested molecules as the compounds properly hydrolyzed by one are generally badly degraded by the other (Table 1). Overall, the multiplicity of possible substrates for BcpE2, and to a lesser extent for BglC, suggests that these enzymes would be part of a “Swiss Army Knife” for providing the bacteria the most readily mobilizable sugar from carbon rich environments. Our results demonstrate that the addition of salicin, one of the best substrates hydrolyzed by BcpE2, as supplement nutrient in the culture medium strongly induces the production of BcpE2 (Figure 3). In *S. venezuelae*, a similar situation has been reported where the production of two distinct β-glucosidases responds to either cellobiose or salicin. The enzyme induced by salicin hydrolyzes aryl β-glucosides but is poorly active on cellobiose while the other enzyme is highly active on – and induced by – cellobiose^31^. Due to their similarity in terms of catalytic specificity and responsiveness to either cellobiose or salicin, we speculate that these two enzymes of *S. venezuelae* are the orthologues of BglC (VNZ_32510) and BcpE2 (VNZ_10820), which share 70%, and 75% of identity with the proteins of *S. scabiei*, respectively. This finding further suggests that this type of GH3 β-glucosidase is not exclusive to *Streptomyces* species that are phytopathogenic However, the orthologue of BcpE2 found in *S. scabiei* is most likely involved in additional functions related to pathogenicity as it is the case for BglC^10^. Indeed, importantly, BcpE2 could also compensate for the loss of a recently discovered function of BglC, i.e., the glucosidase activity on the phytoalexin scopolin^18^. *in vitro* TLC assay revealed that scopolin is the only natural plant compound to be degraded in common by BcpE2 and BglC. Interestingly, the aglycone moiety of scopolin is scopoletin, which has been described as a strong inhibitor of thaxtomin A biosynthesis in *S. scabiei*^19^. Since scopoletin and scopolin have been reported to be produced by plant roots or tubers especially in response to the application of thaxtomin A or under stress conditions such as pathogen infection ^19,27–30^, *S. scabiei* is very likely to encounter these molecules upon host colonization. It is therefore tempting to speculate that BglC and BcpE2 are both involved in the management of these compounds and that BcpE2 would be overproduced in the absence of BglC to take over the contribution of the latter.

**Figure 5.**
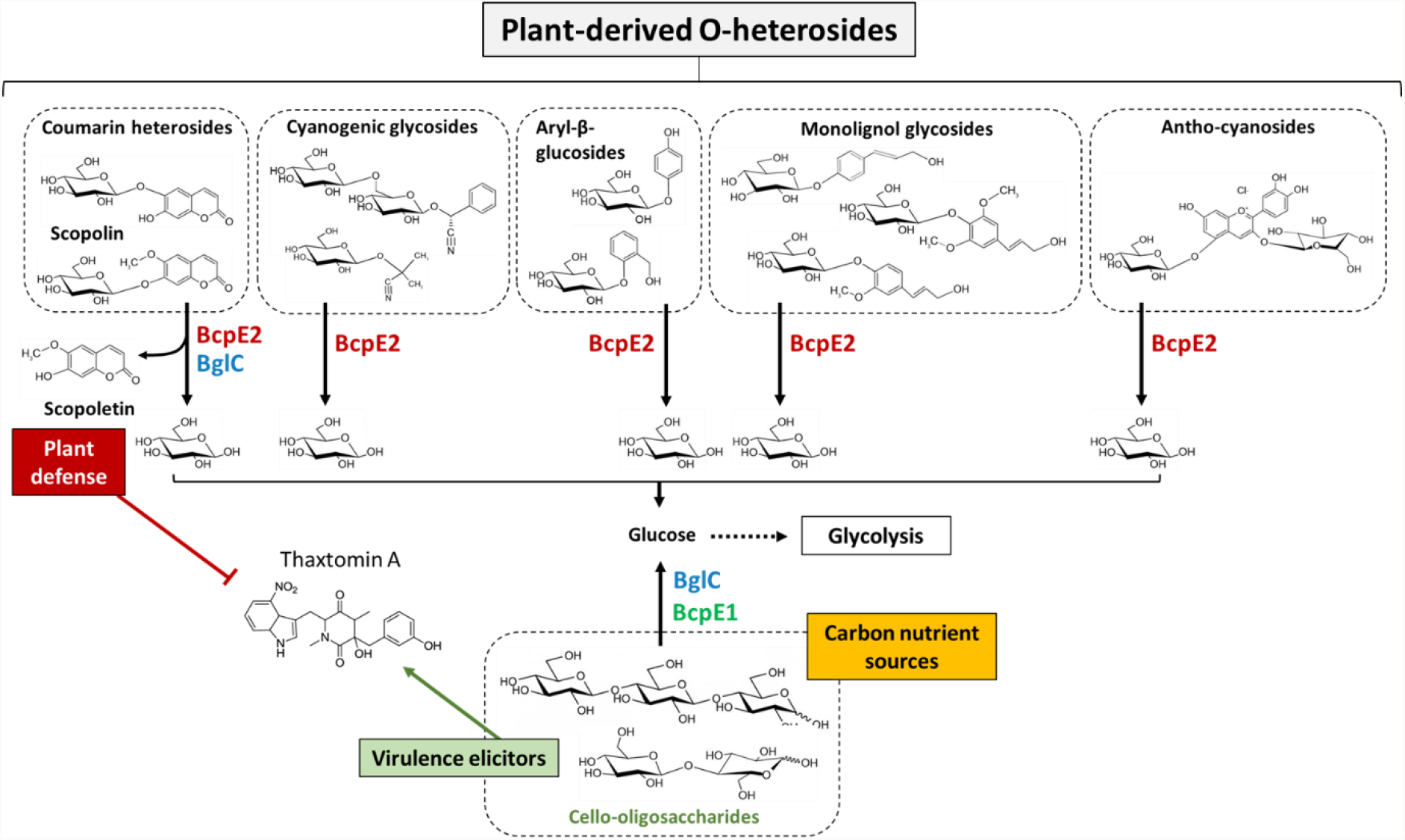
BcpE1 and BcpE2-mediated enzymatic compensation for the loss of BglC. BcpE1 (green) can compensate the activity of BglC (blue) by generating glucose from the hydrolysis of cello-oligosaccharides cellobiose and cellotriose. BcpE2 (red) can also fuel glycolysis by removing glucose from multiple plant heterosides. In addition, BcpE2 can also compensate the role of BglC in plant defense mechanism by displaying a substrate inhibition kinetic profile on scopolin, thereby generating the potent thaxtomin A production inhibitor scopoletin.

### Perspectives

An important question that remains to be answered is “*what are the environmental triggers that induce the expression of bcpE2”*. In other words, “*does the substrate promiscuity of BcpE2 correlate with a mechanism of expression control of bcpE2 sensitive to multiple and dissimilar compounds?”*. We are currently seeking for the transcription factor that controls the expression of *bcpE2* orthologues in streptomycetes. Will this transcription factor be able to sense the presence of multiple substrates of BcpE2 or instead only few structurally similar substrates that would somehow witness for the possible presence of plant-heterosides? The answer to this question is crucial for properly understanding the role of this versatile enzyme. In addition, inactivation of orthologues of *bcpE2* in other model streptomycetes should provide further insight into the importance of this promiscuous enzyme and explain the success of these filamentous bacteria in colonizing plant-derived organic soils.

## Materials and methods

### Strains, chemicals, and culture conditions

Two strains of *Escherichia coli* were used in the present work: (i) DH5α for routine molecular biology applications, and (ii) BL21(DE3) Rosetta™ (Novagen) for heterologous proteins production. Both *E. coli* strains were cultured in LB (BD Difco LB broth) medium supplemented with the appropriate antibiotics (kanamycin (50 µg/mL), chloramphenicol (25 µg/mL)). *Streptomyces scabiei* 87-22 was routinely cultured at 28°C. Tryptic Soy Broth (TSB, Sigma-Aldrich, 30 g/L) was used for liquid pre-cultures. The modified TDM (thaxtomin defined medium ^14^, (Johnson et al., 2007)) minimal medium was prepared as described in (Jourdan et al., 2018) and after autoclaving were supplemented with filter-sterilized carbon sources. The substrates used in this study were purchased from Carbosynth (Cellobiose, Amygdalin, Linamarin, Xylobiose, Laminaribiose, Gentiobiose, Salicin, Arbutin, and Syringin), or from Sigma-Aldrich (4-Nitrophenyl-β-D-glucopyranoside (pNPβG), Esculin, Cyanin chloride, 4-Methylumbelliferyl β-D-glucopyranoside (4-MUG), Coniferin (Abietin), and p-Coumaryl alcohol 4-O-glucoside).

### Heterologous production of His_6_-tagged proteins and purification

BcpE2-His_6_ and His_6_-BglC were produced in *E. coli* BL21(DE3) Rosetta™ transformed with plasmids pBDF004 and pSAJ022, respectively, and purified by nickel affinity chromatography as already described in ^10,20^. The pure proteins were stored at -20 °C and used in HEPES buffer (50 mM, pH 7.5).

### Determination of the pH and temperature optima of BcpE2-His_6_

The β-glucosidase activity was typically determined by the degradation of 4-Nitrophenyl-β-D-glucopyranoside (pNPβG). 95 µL of a determined BcpE2-His_6_ concentration diluted in HEPES buffer (50 mM, pH 7.5) were mixed with 5 µL of pNPβG (20 mM). After incubation at 25°C, the reaction was stopped by the addition of 100 µL of Na_2_CO_3_ (2 M). The release of *para*-nitrophenol was monitored by measuring the absorbance at 405 nm with a TECAN Infinite^®^ 200 PRO. The temperature optimum was determined by varying the incubation temperature from 5 to 60 degrees Celsius with 5°C increments. The pH optimum was determined by varying the pH of the reaction with the use of 3 distinct buffers, *i*.*e*., (i) MES buffer (50 mM) for pH ranging between 5.0 and 6.5, (ii) HEPES buffer (50 mM) for pH ranging between 7.0 and 8.5, and (iii) CHES buffer (50 mM) for pH ranging between 9.0 and 10.0. The measured activity was reported to the maximal value obtained in each experiment which was set to 100%. The results are presented in supplementary Figure S3.

### TLC for hydrolysis of cello-oligosaccharides

Semi-quantitative substrate degradation was assessed by thin layer chromatography (TLC). Reactions were carried out with the BcpE2-His_6_ and His_6_-BglC enzymes (1 µM) and the substrates (5 mM) in HEPES 50 mM pH 7.5 at 40°C for 10 min. At the end of the reaction, the mixture was incubated for 5 min in a boiling water bath to inactivate the enzyme. 1-µL samples of the inactivated reaction mixtures were spotted next to undigested standards on aluminum-backed TLC plates (Silica gel Matrix, Sigma-Aldrich) and thoroughly dried. The protocol, adapted from ^32^, consisted in eluting the loaded TLC plate in a TLC chamber filled with an elution buffer (Chloroform – Methanol – Acetic acid – Water (50:50:15:5 (v/v))). After air-drying the eluted plate, sulfuric acid (5%) in ethanol was sprayed onto the TLC plate and the excess liquid was drained. The revelation was conducted by heating the TLC plate on a hot plate.

### Determination of kinetic parameters for BcpE2-His_6_

The hydrolysis of non-chromogenic substrates for β-glucosidases was followed by glucose quantification, either by HPLC (see HPLC quantification of glucose) or with the D-Glucose HK Assay kit (Megazyme) following the microplate procedure. BcpE2-His_6_ was mixed with the substrate at variable concentrations in HEPES 50 mM pH 7.5, and the incubation was conducted at 40°C for 4 min. The reaction was terminated by a 5-min incubation in a boiling water bath. At least 10 concentrations – if possible distributed around the K_m_ value – were tested in triplicate for each substrate to estimate initial velocity values. The obtained data – initial velocity (V_i_, mM/min) in function of substrate concentration ([S], mM) – were fitted to the Henri-Michaelis-Menten equation V_i_ = (V_max_*[S])/(K_m_ + [S]) using the GraphPad Prism (version 9.2.0) software. K_m_ (mM), V_max_ (mM/min), k_cat_ (s^-1^) and the specificity constant (k_cat_/K_m_ (mM^-1^s^-1^)) were determined for each substrate. Substrate inhibition constants were also determined with GraphPad Prism following the equation V_i_ =(V_max_*[S])/(K_m_ + [S]*(1+[S]/K_i_)).

### HPLC quantification of glucose

Glucose quantification was performed on a Waters HPLC device composed of a Separation Module (e2695) and a Refractive Index (RI) Detector (2414) set at 50°C. 35 µL of glucose-containing samples from terminated reactions were injected on a Aminex HPX-87P (Bio-Rad) column (300 × 7.8 mm) placed in an oven at 80°C. An isocratic flow of milli-Q water was conducted for 20 min at a flow rate of 0.6 mL/min. The RI detector was set on channel 410 and sensitivity 256, and the measurements were expressed in RI Units (RIU). The peak areas associated with glucose (R_t_ = 11.6 min) were integrated and converted into glucose concentrations based on a linear standard curve ranging from 0.1 ng/µL to 400 ng/µL following the equation: y = 644.09x – 3170.4 (y being the Peak area (µV*sec) and x being the glucose amount (ng) / 10 µL injected).

### Crystallization and structure determination of BcpE2-His_6_

BcpE2 was concentrated to 17.4 mg/mL in HEPES 50 mM pH 7.5 and crystallized using the sitting-drop vapor diffusion method. 0.2 µL of protein was mixed with 0.2 µL of precipitant solution (methylpentane-2,4-diol (MPD) 45%, Tris-HCl 0.1M pH 8.5 and 0.2M ammonium acetate) and crystals grew at room temperature. The crystals were transferred into a cryoprotectant solution containing 50% MPD and 50% polyethylene glycol 400 before flash-freezing in a liquid nitrogen bath. Diffraction data were collected at the Soleil Synchrotron Proxima 2a beamline (Paris). Data were integrated and scaled using XDS (X-ray Detector Software ^33^). Initial phases were obtained by molecular replacement using the structure of DesR from *S. venezuelae* as a search model (PDB code 4I3G ^22^) using Phaser^34^. The structure was built with Coot (Crystallographic object-oriented toolkit ^35^) and refined with BUSTER refine^36^). The figures were prepared using PyMOL (The PyMOL Molecular Graphics System, Version 2.4.1 Enhanced for Mac OS X, Schrödinger, LLC.).

### Computational tools

The structure-based alignment was built using the MultAlin-ESPript (3.0) combined tool ^37,38^ using the structure of BcpE2 (PDB code: 7PPJ) as reference for positioning secondary structure elements. The sequences of seven additional characterized GH3 enzymes were included in the alignment (Table S2).

The phylogenetic tree was constructed using the phylogeny.fr tool with the “One Click” mode^39^. The repertoire of characterized bacterial and eukaryotic GH3 proteins was obtained from the CAZy database (accessed on August 27^th^ 2021) and the amino acid sequences obtained were subsequently used in a BLASTp analysis against BcpE2 to select the appropriate proteins for the phylogenetic analysis (Table S2).

The search for PA14 domains was carried out using the MOTIF Search tool (Genome.jp) on the characterized bacterial and eukaryotic GH3 proteins from the CAZy database. The scan for motifs included the Pfam and NCBI-CDD databases in which the pfam07691 or 400161 and 214807 PSSM-Ids were searched for, respectively. In addition, the ScanProsite tool (Expasy) was used on the same amino acid sequences searching for the PA14 (PROSITE entry: PS51820) motif. A manual inspection was conducted to search for the presence of a PA14 domain in the closest GH3s (compared to BcpE2) that were not selected by the search tool. This inspection consisted in a comparison of the predicted secondary structures in the appropriate region of the proteins.

### Determination of the intracellular β-glucosidase activity

Anion exchange chromatography (AXC) to obtain fractions of intracellular β-glucosidases was performed similarly to the method described in ^20^ with protein extracts prepared from cultures of *S. scabiei* 87-22 in TDM supplemented with salicin (0.1%) and/or cellobiose (0.1%). Briefly, 48-hours pre-cultures in TSB were washed twice in TDM without carbon source. After resuspension of the mycelium in the conditions described above, the culture was carried out for 7.5 hours at 28°C. After centrifugation, the mycelium pellet was resuspended in HEPES buffer (50 mM, pH 7.5) and disrupted with an Avestin Emulsiflex C3 homogenizer (3 lysis cycles). The soluble fraction was obtained by centrifugation of the lysed cell suspension and filtering (0.22 µm cut-off) of the supernatant. Using an NGC Quest 10 (Bio-rad) and a HiTrap™ Q HP column (GE healthcare), protein fractions were generated by elution with a linear NaCl gradient (0 to 1 M). The β-glucosidase activity of each fraction was determined by the standard assay using pNPβG as substrate (as described in ^10,20^) and reported to the estimated protein content (Abs_280nm_) of the fraction. These relative activities were then normalized to the maximal activity observed in the second peak (corresponding to BcpE2) of the TDM + cellobiose condition.

### Targeted proteomics analysis

Collection of fractions by anion exchange chromatography and subsequent liquid chromatography – multiple reaction monitoring (LC-MRM) to monitor the relative abundance of BglC and BcpE2 in protein fractions was performed as previously described ^20,40^, and detailed in the supplementary file Table S3.

## Acknowledgements

The work of Be.D. was supported by an aspirant grant from the FNRS (grant 1.A618.18), and a FNRS grant “Crédit de recherche” (grant CDR/OL J.0158.21) to S.R. Ba.D. was supported by a Bijzonder Onderzoeksfonds (BOF, grant 01B08915)-basic equipment from the Ghent University special research funds. F.K. and S.R. are research and senior-research associates of the FRS-FNRS (Brussels, Belgium), respectively. We are very grateful to Sebastien Santini (CNRS/AMU IGS UMR7256) and the PACA Bioinfo platform (supported by IBISA) for the availability and management of the phylogeny.fr website, and to the assistance and support of the team of beamline proxima 2a at the Soleil synchrotron.

## Supplementary data

**Table S1.** Data collection and refinement statistics.

| **BcpE2 (PDB code 7PJJ)** |  |
| --- | --- |
| **Data Collection:** |  |
| Wavelength | 0.98010 |
| Space group | P 3<sub>1</sub> 2 1 |
| a, b, c (Å) | 109.62, 109.62, 164.18 |
| $\alpha$, $\beta$, $\gamma$ (°) | 90, 90, 120 |
| Resolution range (Å)<sup>a</sup> | 47.4 – 3.09 (3.27 – 3.09) |
| Rmerge (%)<sup>a</sup> | 27.3 (222) |
| $\langle I \rangle / \langle \sigma I \rangle$<sup>a</sup> | 6.9 (0.9) |
| Completeness (%)<sup>a</sup> | 99.3 (96.6) |
| Redundancy<sup>a</sup> | 6.7 (6.4) |
| CC 1/2<sup>a</sup> | 0.991 (0.301) |
| **Refinement:** |  |
| Resolution range (Å)<sup>a</sup> | 47.4 – 3.09 (3.2 – 3.09) |
| No. of unique reflections<sup>a</sup> | 21414 (1981) |
| R<sub>work</sub> (%)<sup>a</sup> | 22.3 (33.6) |
| R<sub>free</sub> (%)<sup>a</sup> | 27.0 (35.5) |
| No. Atoms |  |
| Protein | 5904 |
| Solvent | 14 |
| RMS deviations from ideal stereochemistry |  |
| Bond lengths (Å) | 0.009 |
| Bond angles (°) | 1.00 |
| Mean B factor (Å<sup>2</sup>) |  |
| Protein | 90.9 |
| Solvent | 94.3 |
| Ramachandran plot: |  |
| Favored region (%) | 92.7 |
| Allowed regions (%) | 6.4 |
| Outlier regions (%) | 0.9 |

<sup>a</sup> Numbers in parenthesis refer to the highest resolution shell.

### Supplementary Figure S1. Structure-based sequence alignment of BcpE2 and its closest characterized GH3 enzymes

The two catalytic residues – Asp239 and Glu582 – are strictly conserved in all homologous GH3s. The active site of BcpE2 also includes Asp59, Arg127, and three additional aromatic AAs, namely Tyr207, Trp240, and Phe499 which are generally conserved. Asp59 and Arg127 show perfect conservation among all GH3 enzymes displayed in Figure S1, and Trp240 was systematically found next to the catalytic aspartate. Tyr207 was found in all sequences except in BglI of *Schwanniomyces etchellsii* which presented a leucine residue instead. The important Phe499 residue was shared by most GH3-family enzymes considered but sometimes aligned at an adjacent position as in BglI of *S. etchellsii* and KmBglI of *K. marxianus*. Only DesR from *S. venezuelae* and Bgl3B from *Cellulomonas fimi* did not display any phenylalanine residue in this subregion.

**Figure S1.**
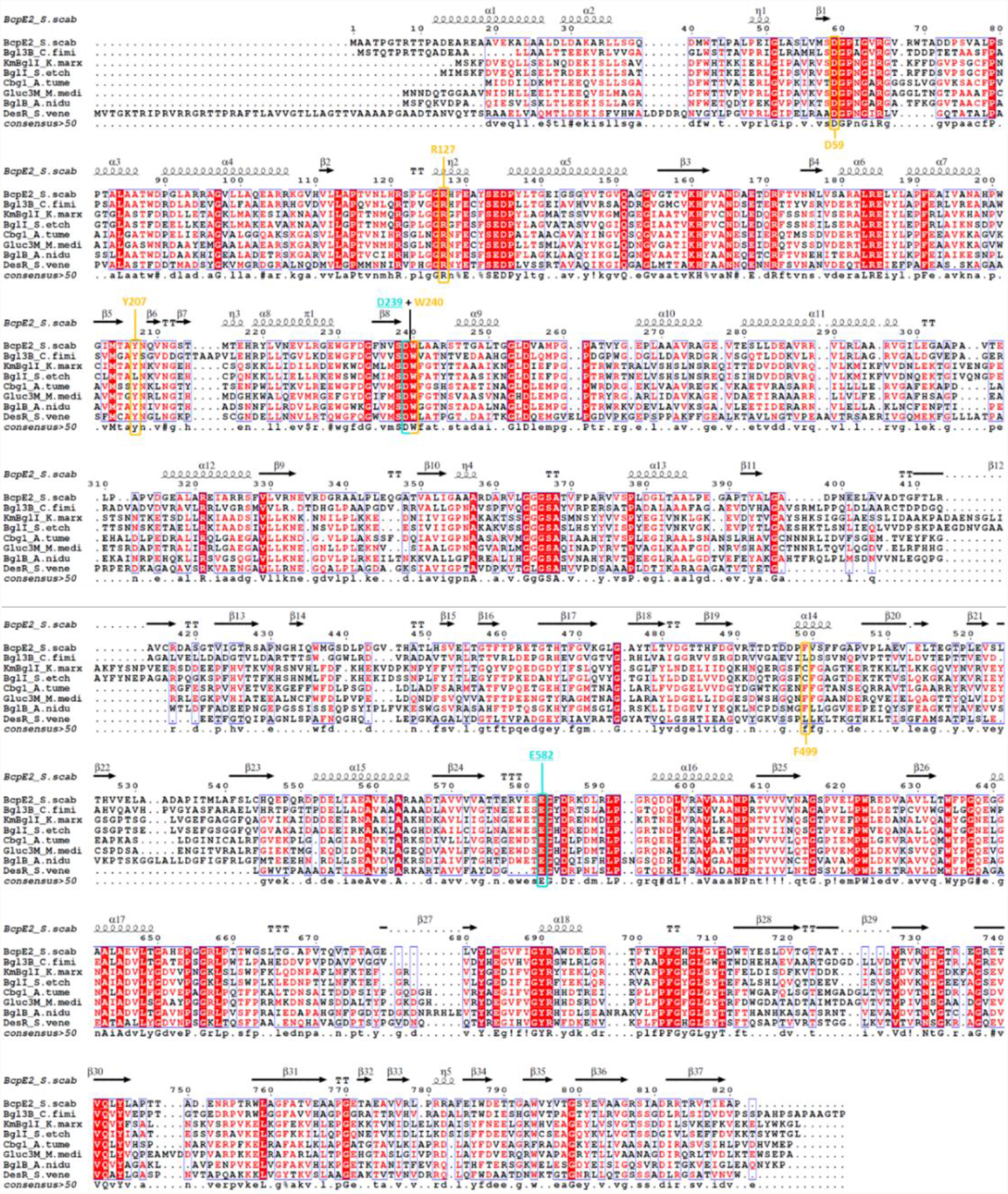
Structure-based sequence alignment of BcpE2 and its closest characterized GH3 enzymes. The residues involved in the active site are indicated in yellow and the catalytic residues, also in the active site, are highlighted in cyan. The red color on residues either indicate perfect conservation (red background) or biochemical similarity (red letter). Blue frames highlight conserved regions for which a consensus motif is found. The “!”, “#”, and “%” symbols in the consensus sequence shows the conservation of branched-chain, acidic or amide, and hydrophobic amino acids (AAs), respectively.

### Supplementary Figure S2. Phylogeny of BcpE2 and its closest characterized GH3-family β-glucosidases

BcpE2 of *S. scabiei* is part of the clade that includes Bgl3B of *Cellulomonas fimi*, Cba from *Cellulomonas biazotea*, Gluc3M of *Martelella mediterranea*, and Cbg1 from *Agrobacterium tumefaciens*. Bgl3B of *Cellulomonas fimi*, and Cba from *Cellulomonas biazotea* are the closest partially characterized non-*Streptomyces* actinobacterial GH3s and have been reported to be active on gentiobiose and cellobiose as natural substrates, respectively (Gao & Wakarchuk, 2014; W. K. R. Wong et al., 1998). BcpE2 also shares a common ancestor with Gluc3M, a cold-active and alkali-stable β-Glucosidase from the Gram-negative rhizobiaceae *Martelella mediterranea*, and with the β-Glucosidase Cbg1 from *Agrobacterium tumefaciens*. Cbg1 is the second closest described homologue though it only has 39% and 55% of AA identity and similarity, respectively, compared to BcpE2. Interestingly, Cbg1 is involved in the virulence induction of some *A. tumefaciens* strains targeting Douglas fir trees (*Pseudotsuga menziesii*) by hydrolyzing the monolignol glucoside coniferin (Castle et al., 1992; Morris & Morris, 1990). Moreover, Cbg1, like BcpE2, is not properly active on cellobiose (Castle et al., 1992; Deflandre et al., 2020) but was instead reported to be active on salicin and arbutin. Beyond the synthetic substrates pNPβG and 4-Nitrophenyl β-D-galactopyranoside (pNPβGal), Gluc3M also exhibited significant activities toward salicin, and konjac powder (glucomannan) (Mao et al., 2010). As Cbg1 hydrolyzes the monolignol glucoside coniferin (Castle et al., 1992), the two other monolignol glucosides syringin and p-coumaryl alcohol 4-O-glucoside were also included as candidate substrates. The coumarin heteroside esculin, which is hydrolyzed by the fungal enzyme -KmBglI GH3 enzyme (Yoshida et al., 2010) - belonging to the phylogenetic clade adjacent to the clade of BcpE2 (Figure S2) was selected as well. The coumarin heteroside scopoline was included because it is a new substrate hydrolyzed by BglC of *S. scabiei* (Deflandre et al. 2022). BcpE2 of *S. scabiei* 87-22 was also suggested by KEGG pathway as candidate beta-glucosidase possibly involved in cyanoamino acid metabolism (https://www.genome.jp/kegg-bin/show_pathway?scb00460+SCAB_64101). Two cyanogenic glucosides were thus also tested as possible targets of BcpE2, *i*.*e*., amygdalin and linamarin. The glycone moiety of amygdalin is the disaccharide gentiobiose and its aglycone part is mandelonitrile, the cyanohydrin of benzaldehyde; linamarin is a glucoside of acetone cyanohydrin. These cyanide-bearing heterosides are plant phytoanticipins whose activation requires the action of a β-glucosidase to release the toxic aglycone moiety from the glycosidic residue. Finally, the natural plant anthocyanidin pigment cyanin, and the synthetic aryl-β-glucoside substrate 4-methylumbelliferyl-β-D-glucoside (4-MUG) were also tested.

**Figure S2.**
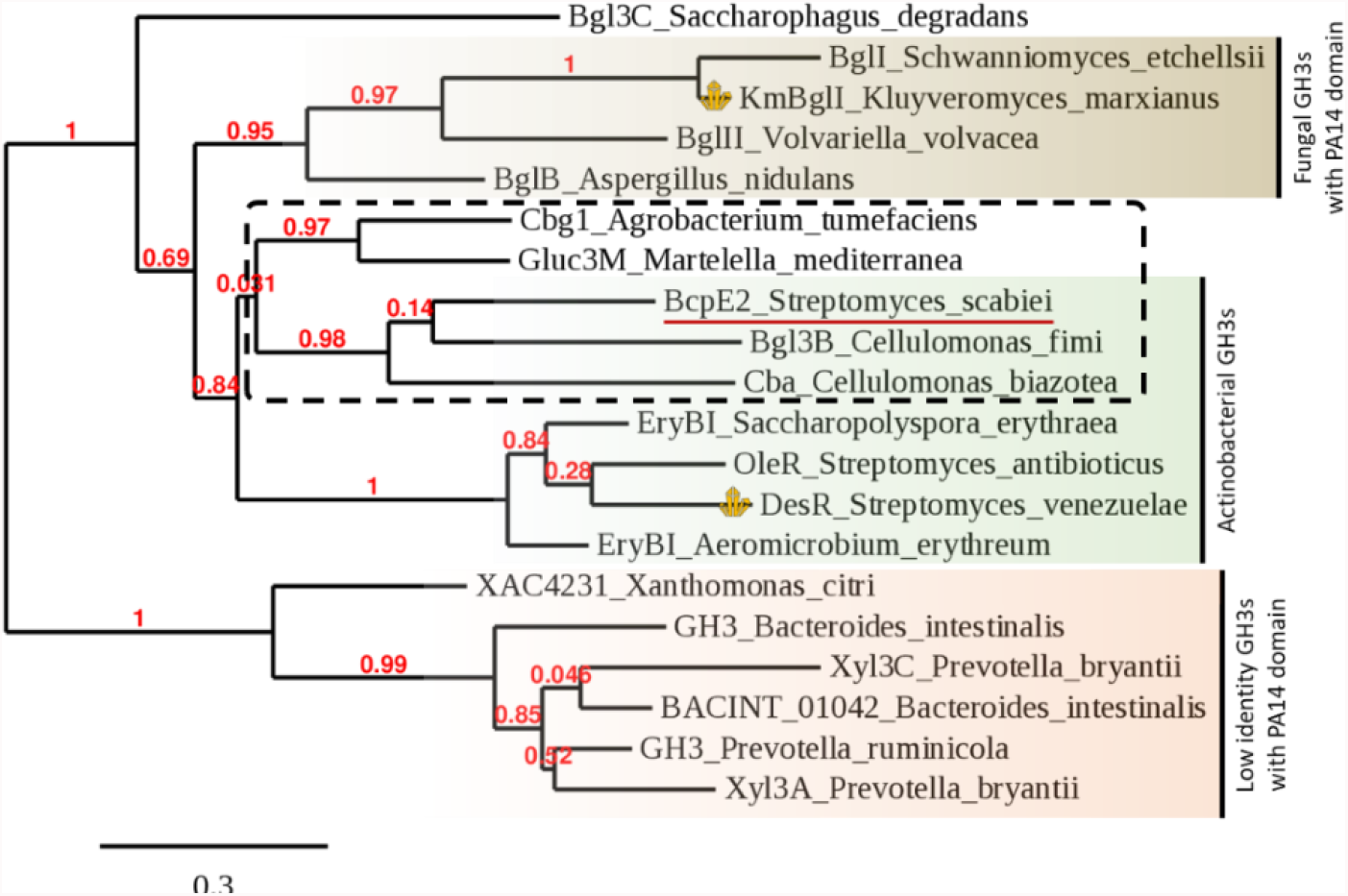
Phylogeny of BcpE2 and its closest characterized GH3-family β-glucosidases in order to identify possible candidate substrates of BcpE2. The 19 closest characterized bacterial and fungal GH3-family beta-glucosidase have been selected based on BLASTp score and/or high query coverage. The dotted square delineates the clade with the closest characterized homologues of BcpE2.

### Supplementary Figure S3. Determination of the pH and temperature optima of BcpE2

The enzyme was obtained as BcpE2-His6 by heterologous production in *E. coli* and subsequent purification by Nickel affinity chromatography as previously described (Deflandre et al., 2020). BcpE2-His6 shows neutral and mesophilic pH and temperature parameters, with optimal activities displayed around pH 6.5-7.5 and 35-40°C, respectively Figure S3). The optimal pH window was relatively narrow, since the enzyme displayed about 60% of its maximal activity at pH values only 0.5 above or below the 6.5-7.5 range (Figure S3, left panel). This pH range is in line with the cytoplasmic compartmentalization of BcpE2 in *S. scabiei*. Below 20°C and above 45°C, the activity rapidly dropped under 50% of the optimal activity (Figure S3, right panel). See the materials and methods section for the detailed protocol.

**Figure S3.**
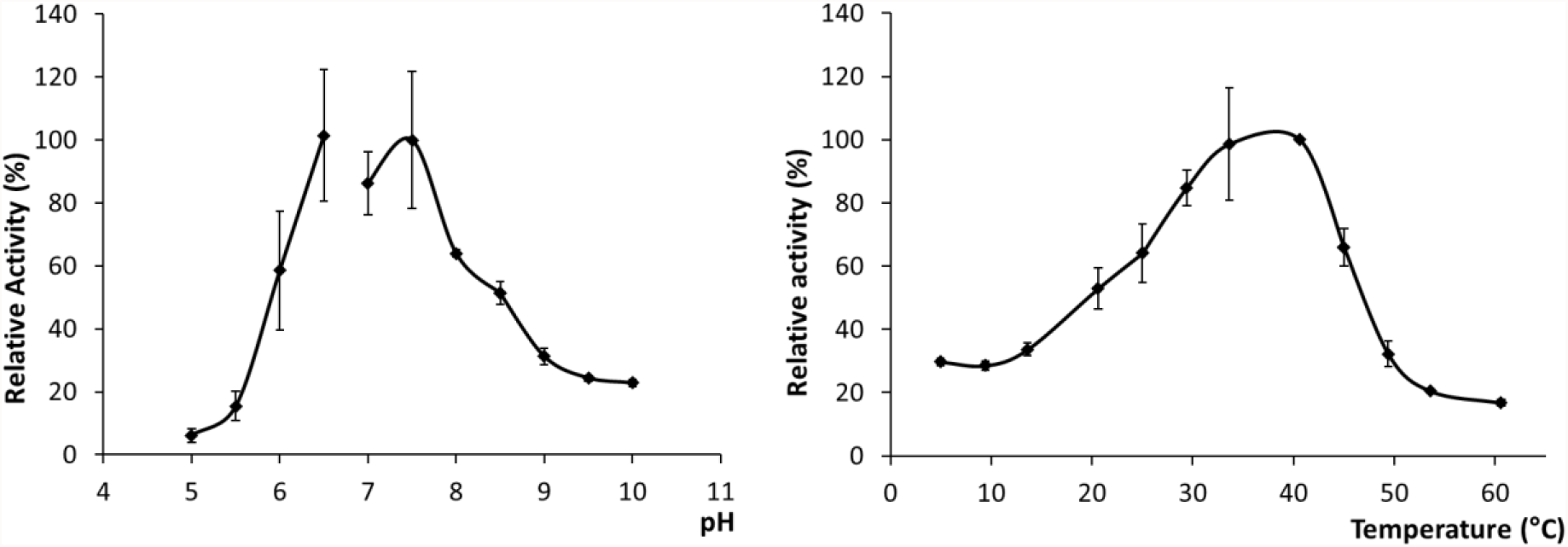
Determination of the pH and temperature optima of BcpE2. Relative activity assays with pNPβG as substrate, normalized to the maximal value measured in each assay. The influence of the pH (left panel) was assessed by increments of 0.5 from pH 5 to pH 10, and the temperature (right panel) was assessed by increments of 5°C from 5 to 60°C.

**Table S2.**
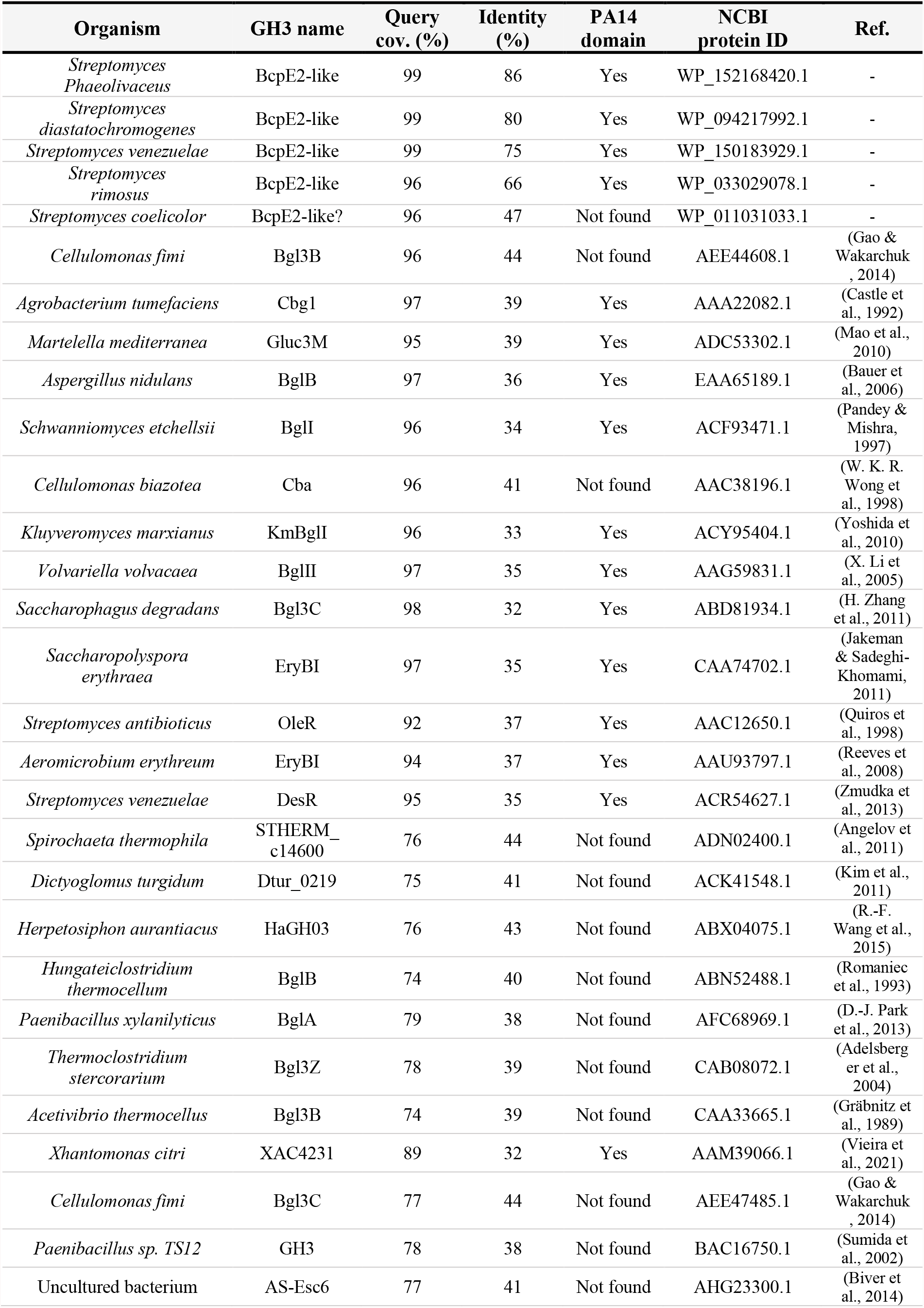
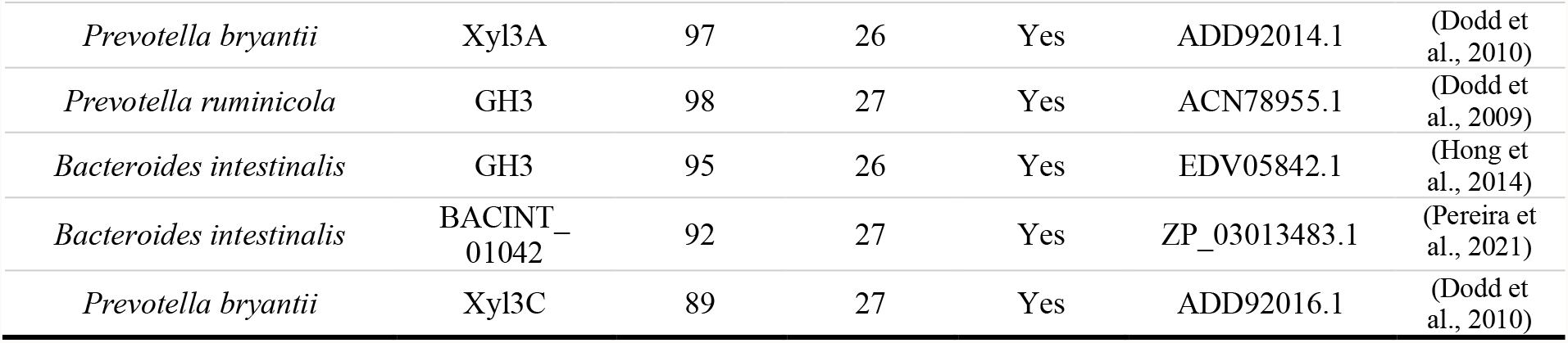
Characteristics of GH3-family proteins homologous to BcpE2 (used for structure-based alignment and phylogenetic tree construction)

**Supplementary Table S3.** Transitions selected for MRM relative quantification of BglC/BcpE2 β-glucosidases.

| Peptide | Precursor ion mass (m/z) | CE (V) | Fragment y-ion mass (m/z) |
| --- | --- | --- | --- |
| BglC |  |  |  |
| LVDELLAK | 450.7737<sup>++</sup> | 16 | 787.456<sup>+</sup><br>688.3876<sup>+</sup><br>573.3606<sup>+</sup> |
| TDPVASLR | 429.7376<sup>++</sup> | 15 | 642.3933<sup>+</sup><br>545.3406<sup>+</sup><br>446.2722<sup>+</sup> |
| BcpE2 |  |  |  |
| AGVLLAQEAR | 514.2984<sup>++</sup> | 18 | 800.4625<sup>+</sup><br>687.3784<sup>+</sup><br>574.2944<sup>+</sup> |
| DASGTVIGTR | 488.7565<sup>++</sup> | 17 | 703.4097<sup>+</sup><br>646.3883<sup>+</sup><br>545.3406<sup>+</sup> |
| AADTAVVVVATTER | 701.8805<sup>++</sup> | 25 | 973.5677<sup>+</sup><br>874.4993<sup>+</sup><br>775.4308<sup>+</sup> |
| BSA |  |  |  |
| AEFVEVTK | 461.7477<sup>++</sup> | 16 | 722.4083<sup>+</sup><br>575.3399<sup>+</sup><br>476.2715<sup>+</sup> |
| QTALVELLK | 507.8133<sup>++</sup> | 18 | 785.5131<sup>+</sup><br>714.476<sup>+</sup><br>601.3919<sup>+</sup> |
| Phos B |  |  |  |
| VFADYEEYVK | 631.8006<sup>++</sup> | 22 | 945.42<sup>+</sup><br>830.3931<sup>+</sup><br>667.3297<sup>+</sup> |
| LLSYVDDEAFIR | 720.8721<sup>++</sup> | 26 | 964.4734<sup>+</sup><br>865.405<sup>+</sup><br>750.3781<sup>+</sup> |
| VLYPNDNFFEGK | 721.8512<sup>++</sup> | 26 | 1067.479<sup>+</sup><br>970.4265<sup>+</sup><br>856.3836<sup>+</sup><br>741.3566<sup>+</sup> |

### Detailed protocol of the targeted proteomic approach

Fractions collected from the anion exchange chromatography (AXC) were subjected to trichloroacetic acid (TCA) precipitation. The AXC fractions were mixed with 100% (w/v) TCA solution (Sigma-Aldrich) (4:1), the proteins were precipitated overnight at 4°C and collected by centrifugation (16,000g; 30 min, 4°C). The protein pellet was washed twice with ice-cold acetone and solubilized in 50 mM ammonium bicarbonate containing 2 M urea. Protein concentrations were estimated by Bradford’s method (Coomassie Plus Protein Assay kit, Pierce). Dried protein (10 µg) was solubilized in 2 M urea/50mM NH4HCO3 spiked with bovine serum albumin (MS-grade protein standard) (1:250) (Thermo-Scientific), and denatured by heating to 80°C for 10 min. The solution was subsequently reduced with 5 mM dithiothreitol (Sigma-Aldrich) for 10 min at 60°C and alkylated with 15 mM iodoacetamide for 20 min at RT in the dark before digestion with trypsin (Promega, Madison, USA) overnight at 37°C (1:50 w/w). Acidified digested samples were desalted using OMIX C18 pipette tips (Agilent). The desalted peptides were dried under vacuum and dissolved in 0.1% formic acid, 3% ACN and 10 fmol/µl phosphorylase B (Hi3 Phos B Standard, Waters).

For BglC and BcpE2 identification by MRM, samples (0.5 µg) were injected onto an ultraperformance liquid chromatography (UPLC) M-class system (Waters) and trapped on a 300 µm x 50 mm, 5 µm, 100 Å Acquity UPLC M-Class Symmetry C18 Trap Colum (Waters). The washing step on the trap column was performed for 2 min with 3% B at a flow rate of 15 µL/min, with solvent A 0.1% HCOOH in H2O (Biosolve) and solvent B 0.1% HCOOH in ACN (Biosolve). Subsequently, the peptides were separated on a 150 µm x 100 mm, 1.8 µm HSS T3, iKey separation device in 10 min at a flow rate of 2 µL/min using a linear solvent B gradient (3-50%). The separated peptides were introduced into the IonKey source coupled to a Waters Xevo TQ-S triple-quadrupole mass spectrometer for detection of the analytes in the positive-ion mode (ESI+). The MRM mode with transitions of selected proteotypic peptides at a set cone voltage and different collision energies for each precursor, was used for detection, normalization (BSA) and MS performance check (Phos B) (Figure S3). Capillary voltage was set at 3.5kV, the cone voltage at 35V and the source temperature at 120°C. In the collision cell, argon was introduced at a flow rate of 0.15 mL/min. Data were acquired with the developed MRM mode (MassLynx 4.1), subsequently uploaded into Skyline (Pino et al., 2020) for data analysis, and, subjected to a Savitsky-Golay Smoothing, the total area under the curve (AUC) for each peptide was calculated and normalized to BSA.

